# Interferon restores replication fork stability and cell viability in BRCA-defective cells via ISG15

**DOI:** 10.1101/2023.03.16.533020

**Authors:** Uddipta Biswas, Ramona N. Moro, Suhas S. Kharat, Prosun Das, Arnab Ray Chaudhuri, Shyam K. Sharan, Lorenza Penengo

**Affiliations:** University of Zurich, Institute of Molecular Cancer Research, 8057 Zurich, Switzerland; Mouse Cancer Genetics Program, Center for Cancer Research, National Cancer Institute, National Institutes of Health, Frederick, 21702 Maryland, USA; Department of Molecular Genetics, Erasmus MC Cancer Institute, Erasmus University Medical Center, 3015GD Rotterdam, the Netherlands

## Abstract

DNA replication and repair defects or genotoxic treatments trigger interferon (IFN)-mediated inflammatory responses. However, whether and how IFN signaling in turn impacts the DNA replication process has remained elusive. Here we show that IFN promotes replication fork stability, cell proliferation and survival in BRCA1/2-defective cancer cells and rescues the lethality of BRCA2-deficient mouse embryonic stem cells. Although IFN activates hundreds of genes, these effects are specifically mediated by the ubiquitin-like modifier ISG15 (IFN-stimulated gene 15). Inactivation of ISG15 or of the enzymes promoting its conjugation, referred as ISGylation, completely suppresses the impact of IFN on the replication process. Depletion of ISG15 significantly reduces cell proliferation rates whereas its upregulation results in increased resistance to the chemotherapeutic drug cisplatin in human BRCA1-mutated triple-negative and mouse BRCA2-deficient breast cancer cells, respectively. Accordingly, cells carrying BRCA1/2 defects consistently show increased ISG15 levels, representing a novel, in-built mechanism of drug resistance linked to BRCAness.

## Introduction

The fidelity of genome duplication is crucial for genome maintenance. A variety of stresses can challenge this fundamental process resulting in ‘replication stress’, characterized by alteration of the rate and the fidelity of DNA synthesis. If not timely and properly addressed, replication stress can lead to replication fork collapse and DNA double-strand breaks, promoting cancer development. Multiple mechanisms and factors regulating DNA replication fork progression and stability cooperate to ensure genome integrity upon replication stress. In fact, germline or acquired mutations targeting these factors represent a relevant percentage of alterations observed in human malignancies. Among others, mutations in *BRCA1/2* genes are associated with an increased risk of developing different types of tumors including breast, ovarian, pancreatic and prostate cancer (Cavanagh and Rogers, 2015). BRCA1 and BRCA2 are key proteins in repairing DNA breaks, by promoting the homologous recombination (HR) DNA repair pathway, and in maintaining the stability of newly synthetized DNA strands, by protecting stalled replication forks from degradation, hence preventing chromosomal aberrations (Schlacher et al., 2011, 2012; Scully et al., 2019).

A side effect of genetic defects or genotoxic treatments challenging replication fork stability is the generation of byproducts (ssDNA, oligonucleotides, DNA:RNA hybrids) and their accumulation in the cytosol. These nucleic acids mimic pathogen infection and are typical activators of GMP–AMP synthase (cGAS), a DNA sensor that triggers innate immune responses through the production of the second messenger cyclic GMP-AMP (cGAMP) and activation of the adaptor protein STING (Härtlova et al., 2015). Downstream activation of TBK1 and IFN regulatory factor 3 (IRF3) then occurs, resulting in the activation of type I IFN and the upregulation of the IFN-stimulated genes, ISGs (Li and Chen, 2018; MacKenzie et al., 2017; Parkes et al., 2017). This inflammatory response, induced by chromosomal instability, has broad and diverse effects depending on the cancer type and context, as acute inflammation have immune-stimulatory effects against malignancies while chronic inflammation may promote cancer development (Bakhoum et al., 2018; Erdal et al., 2017; Harding et al., 2017; MacKenzie et al., 2017), and can even be exploited for immunotherapeutic purposes (Cybulla and Vindigni, 2022), underlining the importance of better understanding the links between replications stress and the immune system.

The mechanisms through which genomic instability triggers the immune response have been extensively investigated in recent years. Yet, the consequences of this activation on DNA replication, cell fitness and homeostasis are far less understood. We recently reported that type I IFN (i.e., IFNβ), through the up-regulation of ISG15, promotes deregulated DNA replication fork progression, representing the first natural event leading to an acceleration of DNA replication rate, with detrimental consequences for the cells (Raso et al., 2020). ISG15 is a ubiquitin-like modifier that exerts its functions via covalent conjugation to targets, via non-covalent interaction and as secreted molecule (Dos Santos and Mansur, 2017; Swaim et al., 2017; Villarroya-Beltri et al., 2017). ISG15 plays a central role in the antimicrobial response by protecting the host during infection (Perng and Lenschow, 2018), but it is also frequently deregulated in cancer (Han et al., 2018), yet its exact role is controversial. Recently, ISG15 (and its conjugating enzymes) has been reported to modulate DNA replication and genome stability (Park et al., 2014; Raso et al., 2020; Wardlaw and Petrini, 2022), though mechanisms and functions have remained elusive. Here, we addressed the effects of IFNβ signaling – and ISG15 – on pathological genetic contexts, the BRCA1/2-deficiencies, characterized by high genomic instability, DNA repair defects and severe replication stress. We found that treatment with low doses of IFNβ completely restores replication fork stability in BRCA1/2-deficient cells. This effect, observed invariantly across multiple human and mouse cell lines, including patient-derived BRCA1/2 defective lines, is entirely dependent on ISG15 and on the enzymes mediating its covalent conjugation – referred as ISGylation – namely UBE1L and TRIM25. Remarkably, IFNβ treatment is also able to rescue the viability of BRCA2-deficient mouse embryonic stem cells (mESCs) in an ISG15/ISGylation-dependent manner and the up-regulation of ISG15 confers drug resistance to BRCA2-deficient cells. Consistent with this, loss of ISG15 has dramatic effects on cell fitness, impairing proliferation of BRCA1-mutated triple-negative breast cancer cells. Altogether, these findings reveal that the IFN/ISG15 system controls fork stability in clinically relevant pathological contexts, increasing the fitness of BRCA1/2-deficient cancer cells and affecting the drug response.

## Results

### IFNβ fully restores the stability of stalled replication forks in BRCA1/2-deficient cells

Acute type I IFN treatment is highly toxic for cells, including cancer cells; yet the effect of chronic, low dose IFN treatment has been poorly explored. We recently reported that low doses of IFNβ treatment accelerates DNA synthesis in many different cell types, by increasing DNA replication fork speed (Raso et al., 2020). We have now examined the effects of type I IFN signaling on BRCA1/2-deficient cells, which are characterized by extensive degradation of newly synthetized DNA strand upon fork stalling, impaired DNA repair pathway by HR and – at least initially – exquisite sensitivity to genotoxic chemotherapeutic treatments.

To monitor the effect of IFNβ treatment on the stability of nascent DNA strands in BRCA1/2-deficient cells, we took advantage of the DNA fiber technique, in which ongoing DNA synthesis is labelled with halogenated thymidine analogues (CldU followed by IdU), which can be recognized by specific antibodies. To measure the stability of stalled replication forks, after CldU/IdU labelling cells are treated with hydroxyurea (HU) to deplete the cellular pool of dNTPs and induce fork pausing. In control cells, the arrested forks are protected by several factors, including BRCA1, BRCA2, RAD51, FANCA, FANCD2 (Rickman and Smogorzewska, 2019), and therefore the ratio between IdU (green) and CldU (red) is approximately 1 (see scheme in Figure 1a). As expected (Schlacher et al., 2011), cells depleted of BRCA2 (siBRCA2) show excessive degradation of the newly synthetized DNA (green tract) compared to control cells (siLuc), leading to a IdU/CldU ratio around 0.6 (Figure 1b–1d). Remarkably, pre-treatment of cells with IFNβ completely restores the stability of nascent DNA (siBRCA2 + IFNβ). A similar effect was also observed in BRCA1-depleted cells (Figure 1e–1g), indicating a general effect of IFNβ treatment in promoting the stability of newly synthetized DNA in BRCA1/2-deficient cells. To extend our investigations into a more clinically relevant context, we tested the effect of IFNβ treatment on pancreatic adenocarcinoma cells (CAPAN-1) and triple-negative breast cancer cells (SUM149PT and MDA-MB-436) carrying BRCA1 or BRCA2 defects and we invariably obtained the same result (Figure 1h–1j).

**Figure 1.**
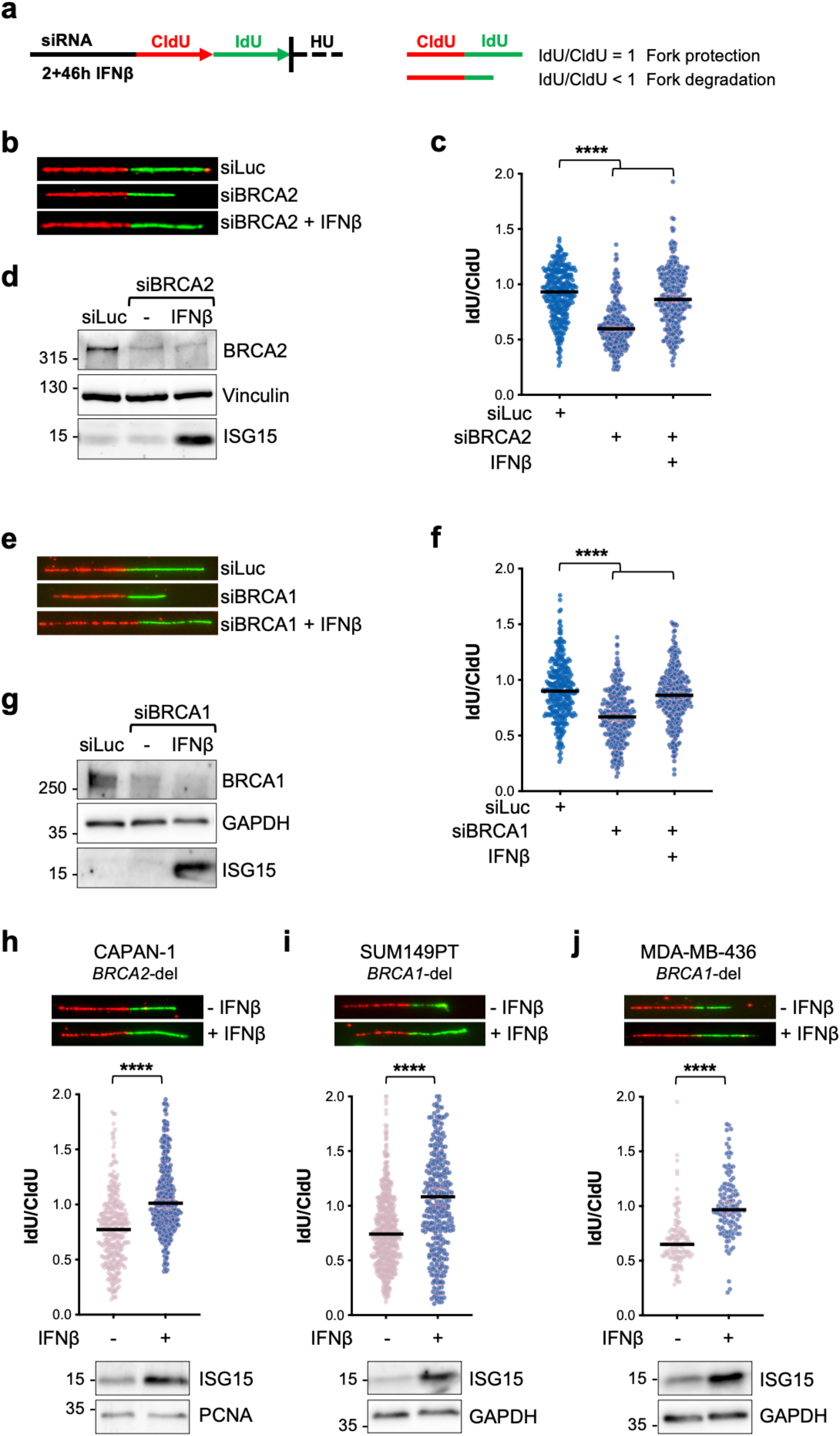
IFNβ treatment promotes DNA replication fork protection in BRCA1/2 depleted cells. (**a**) Schematic diagram of DNA replication fork degradation assay. Cells were transfected with siRNAs and optionally treated with IFNβ (30 U/mL, 2 h + 46 h release), followed by incubation with CldU (red) and IdU (green) and treatment with hydroxyurea (HU, 4 mM) for 4 h. IdU/CldU ratios < 1 indicate fork degradation. (**b**) Representative images of fibers scored in siBRCA2 or siLuc U2OS cells treated with ± IFNβ (30 U/mL, 2 h) and chased for 46 h. (**c**) IdU/CldU ratio analysis in siBRCA2 or siLuc U2OS treated with ± IFNβ (30 U/mL, 2 h) and chased for 46 h (n=3). Horizontal lines represent the median value. Statistical analysis according to Kruskal-Wallis test was performed; ****, P<0.0001. (**d**) ISG15 and BRCA2 protein expression as in **c**. Vinculin shows equal loading and ISG15 induction reveals activation of IFNβ pathway. (**e**) Representative images of fibers scored in siBRCA1 or siLuc U2OS cells treated with ± IFNβ (30 U/mL, 2 h) and chased for 46 h. (**f**) IdU/CldU ratio analysis in siBRCA1 or siLuc U2OS cells treated with ± IFNβ (30 U/mL, 2 h) and chased for 46 h (n=3). Horizontal lines represent the median value. Statistical analysis according to Kruskal-Wallis test was performed; ****, P<0.0001. (**g**) ISG15 and BRCA1 protein expression as in **f**. (H-J) IdU/CldU ratio analysis in CAPAN-1 (H), SUM149PT (I) and MDA-MB-436 (J) cells along with ISG15 protein expression with ± IFNβ (30 U/mL, 2 h) treatment and chased for 46 h (n=3). Horizontal lines represent the median value. Statistical analysis according to Mann-Whitney test was performed; ****, P<0.0001.

### ISG15 is required and sufficient for IFNβ-induced fork protection

To explore the molecular determinants of IFNβ-mediated fork protection in BRCA2-deficient cells, we first focused on ISG15, a ubiquitin-like protein strongly induced by type I IFN that we recently reported to play a key role in modulating replication fork progression (Raso et al., 2020). Using a similar experimental setting as in Figure 1, we tested the effect of ISG15 loss on IFNβ-mediated fork protection in BRCA2-deficient cells. Remarkably, depletion of ISG15 completely reversed the effect of IFNβ on fork stability, clearly revealing the essential role of ISG15 in IFNβ-mediated stalled fork protection (Figure 2a and 2b). Furthermore, we extended these analyses to mouse embryonic fibroblasts (MEFs) and confirmed that loss of BRCA2 results in extensive degradation of stalled replication forks similarly to human cells (Figure 2c). Notably, fork degradation in these MEFs is reversed by the treatment with mouse IFNβ and is completely dependent on ISG15 expression, indicating an evolutionary conserved function of IFN and ISG15 in the replication process (Figure 2c, S1a and S1b).

**Figure 2.**
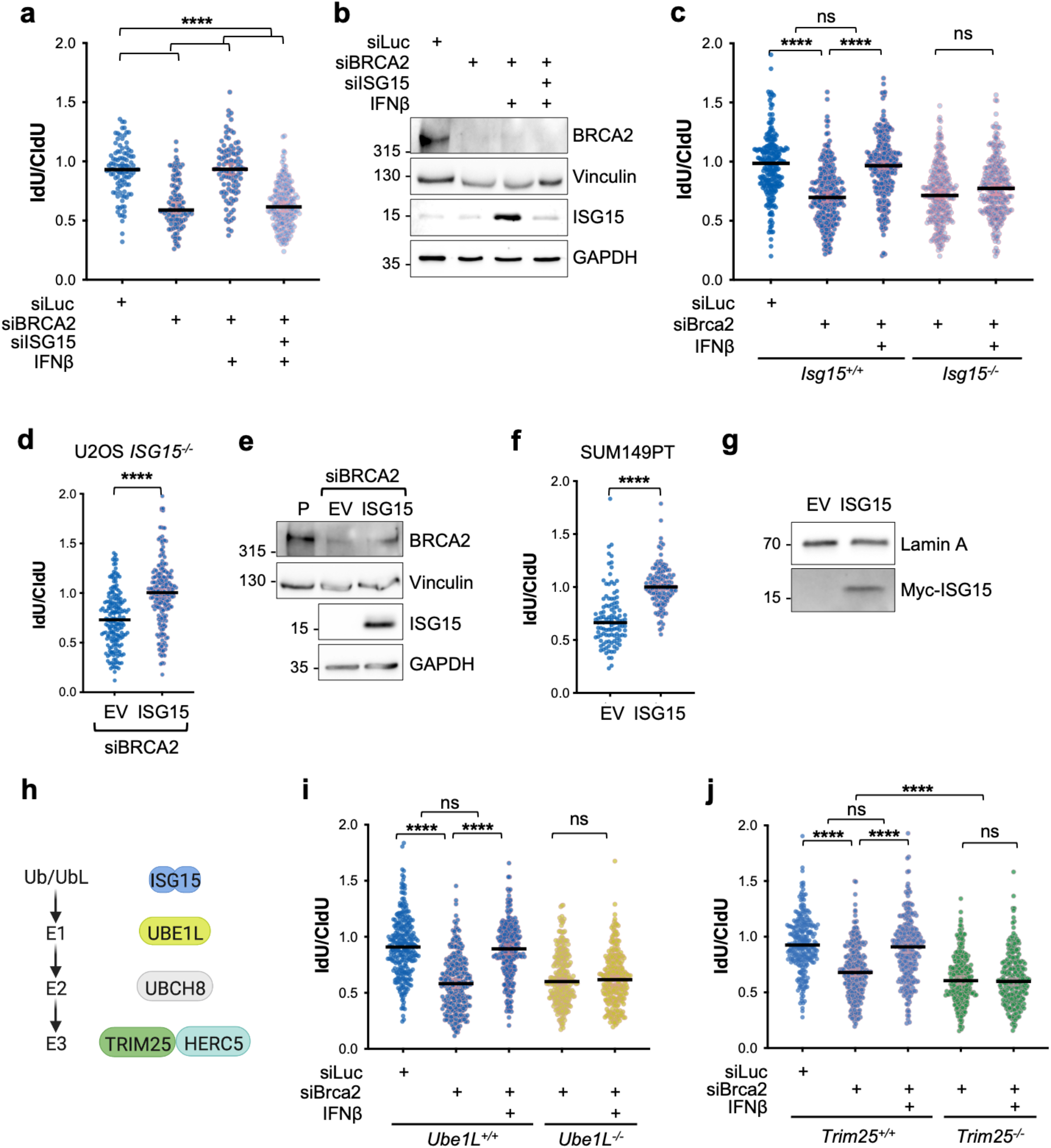
IFN/ISG15 restores fork protection in BRCA-deficient cells via ISG15 conjugation. (**a**) IdU/CldU ratio analysis in BRCA2 or ISG15 depleted U2OS cells treated with IFNβ (30 U/mL, 2 h) and chased for 46 h (n=2). Horizontal lines represent the median value. Statistical analysis according to Kruskal-Wallis test was performed; ****, P<0.0001. (**b**) ISG15 and BRCA2 protein expression as in **a**. (**c**) IdU/CldU ratio analysis in parental (*Isg15*^+/+^) and *Isg15*^-/-^ MEF cells transiently depleted of BRCA2 (siBrca2) and treated with ± IFNβ (30 U/mL, 2 h) and chased for 46 h (n=2). Horizontal lines represent the median value. Statistical analysis according to Kruskal-Wallis test was performed; ns, non-significant; ****, P<0.0001. (**d**) IdU/CldU ratio analysis in BRCA2 depleted *ISG15*^-/-^ U2OS cells expressing empty vector (EV) or FLAG-ISG15 (n=2). Horizontal lines represent the median value. Statistical analysis according to Mann-Whitney test was performed; ****, P<0.0001. (**e**) ISG15 and BRCA2 protein expression as in **d**. P, parental U2OS. (**f**) IdU/CldU ratio analysis in SUM149PT (*BRCA1*-deleted) cells expressing empty vector (EV) or Myc-ISG15 (n=3). Horizontal lines represent the median value. Statistical analysis according to Mann-Whitney test was performed; ****, P<0.0001 (**g**) Myc-ISG15 staining as in **f**. (**h**) Schematic of the machinery involved in ISG15 conjugation. (**i**) IdU/CldU ratio analysis in parental (*Ube1L*^+/+^) and *Ube1L*^-/-^ MEF cells transiently depleted of BRCA2 (siBrca2) and treated with ± IFNβ (30 U/mL, 2 h) and chased for 46 h (n=3). Horizontal lines represent the median value. Statistical analysis according to Kruskal-Wallis test was performed; ns, non-significant; ****, P<0.0001. (**j**) IdU/CldU ratio analysis in parental (*Trim25^+/+^*) and *Trim25*^-/-^ MEF cells transiently depleted of BRCA2 (siBrca2) and treated with ± IFNβ (30 U/mL, 2 h) and chased for 46 h (n=3). Horizontal lines represent the median value. Statistical analysis according to Kruskal-Wallis test was performed; ns, non-significant; ****, P<0.0001.

We next addressed whether the sole upregulation of ISG15 is sufficient to induce this phenotype, in absence of IFNβ stimulation. We used the ISG15 knockout (*ISG15*^-/-^) Flp-In TREx U2OS cells that we previously described (Raso et al., 2020) and found that the doxycycline inducible expression of FLAG-ISG15 can restore fork protection in BRCA2-deficient cells (Figure 2d and 2e). Moreover, we engineered SUM149PT cells, by lentiviral transduction, to stably integrate a plasmid expressing Myc-ISG15 upon doxycycline treatment and confirmed that the sole expression of ISG15 restores fork stability in BRCA1-defective contexts (Figure 2f and 2g).

### The ISG15 conjugation machinery is required for IFNβ-induced fork protection

Being part of the ubiquitin family, an important mechanism of action of ISG15 is via covalent conjugation to target proteins by means of the E1 activating, E2 conjugating and E3 ligating enzymes (Figure 2h). Hence, we investigated whether the enzymes involved in ISGylation are required for the restoration of fork stability in BRCA2-deficient cells. We first tested the effect of genetic ablation of the ISG15-specific E1 enzyme (UBE1L) in MEFs (*Ube1L*^-/-^) and found that it is strictly required for the IFNβ-dependent fork protection in BRCA2-deficient cells (Figure 2i, S1c and S1d). The two major ISG15 ligases promoting ISGylation are HERC5 and TRIM25 (Wong et al., 2006; Zou and Zhang, 2006). Since HERC5 is mainly associated to innate antiviral response and has been reported to be rather promiscuous in terms of target specificity, we first focused on TRIM25 and found that loss of TRIM25 in MEFs (*Trim25*^-/-^) prevented restoration of replication fork stability upon IFNβ stimulation (Figure 2j, S1e and S1f). Similar results were obtained in U2OS cells where IFNβ-dependent fork stabilization in BRCA2-deficient cells is reversed by loss of ISG15, UBE1L and TRIM25 (Figure S1g–S1i). Conversely, we observed no contribution of HERC5 in IFN-mediated fork protection in BRCA2-depleted U2OS cells (data not shown). Altogether these results show that ISG15 conjugation is required for IFNβ-mediated fork protection in BRCA2-deficient context.

### IFNβ does not rescue RAD51 *foci* in BRCA1/2-deficient cancer cells

BRCA1 and BRCA2 are important to promote homology-directed DNA repair, by promoting the loading of RAD51 onto chromatin, a key step in the HR. Consequently, RAD51 accumulation in discrete nuclear *foci* following DNA damage is largely impaired in BRCA1/2-deficient cells. To better assess the effect of the IFN/ISG15 system in these genetic contexts, we tested whether its upregulation exerts any effect on the restoration of chromatin loading of RAD51. Thus, we monitored the formation of RAD51 *foci* upon etoposide treatment (ETO) in BRCA1-defective cells (SUM149PT) and observed that the accumulation of RAD51 *foci* was highly impaired in ETO-treated cells and IFNβ treatment did not show any effect on the localization of RAD51 (Figure S2a–S2c). We obtained analogous results using the BRCA2-deficient (*Brca2*^-/-^) mouse mammary tumor cell line KB2P (clone 1.21(Evers et al., 2008)), confirming that treatment with IFNβ did not restore RAD51 *foci* (Figure S2d–S2f). Next, we performed similar experiments in U2OS cells upon transient depletion of BRCA1 or BRCA2. RAD51 *foci* induced by ETO treatment and by ionizing radiation (IR) were readily detectable in control cells (siLuc) and, expectedly, highly reduced both in BRCA1- and BRCA2-depleted cells. Here, IFNβ treatment partially restored RAD51 *foci* as compared to BRCA1/2-proficient cells (Figure S2g–S2l), probably reflecting the residual amount of BRCA1 and BRCA2 still present in the cells.

### IFNβ restores the viability of BRCA2-deficient mESCs via ISG15 conjugation

*BRCA1* and *BRCA2* genes are essential in mammals, and their knockout leads to early embryonic lethality in mice (Gowen et al., 1996; Hakem et al., 1996; Sharan et al., 1997). It has been reported that the survival of *Brca2*-deficient mESCs can be promoted by restoration of fork protection, while HR is still impaired (Ray Chaudhuri et al., 2016). Hence, we asked whether IFNβ/ISG15, by promoting replication fork protection in BRCA2-deficient cells, can also restore viability of ESCs. To address this, we used PL2F7 mESCs that carry one functionally null and one conditional allele of *Brca2* (*Brca2*^*flox*-/-^; Figure 3a and S3a). Here, the transfection with CRE recombinase generates a functional HPRT minigene, which in principle allows these cells to grow in HAT (hypoxanthine, aminopterin and thymidine) medium, and promotes the deletion of the conditional allele, leading to complete loss of *Brca2* and thereby to lethality (Ding et al., 2016; Kuznetsov et al., 2008). To test the effect of IFNβ on the viability of mESCs, we optionally treated these cells with mouse IFNβ, either for 48 h in continuous or for 2 h pulse and 46 h chase, prior to CRE transfection and selection in HAT medium (Figure 3b and S3b). While the genotyping of the very few surviving colonies did not reveal any *Brca2*^-/-^ clones in untreated cells (DMSO), confirming the essential role of BRCA2 in ESCs, pretreatment with IFNβ led to a remarkable number of viable *Brca2*^-/-^ clones, exceeding 30%, indicating that the upregulation of IFNβ signaling efficiently rescues the viability of *Brca2*-deficient mESCs (Figure 3c and 3d). Consistently, also in this system we observed rescue of replication fork protection following IFNβ, although to a lesser extent compared to other systems (Figure S3c), without restoration of RAD51 *foci* (data not shown). To investigate the possible role of the ISG15 system in this context, we monitored the effect of loss of ISG15 and of the ISGylating enzymes UBE1L and TRIM25 on the IFNβ-induced on viability of *Brca2*^-/-^ cells. Remarkably, even upon IFNβ treatment, we obtained no clones that are *Brca2*^-/-^ in cells depleted of ISG15, UBE1L and TRIM25 (Figure 3e, 3f, S3d and S3e), highlighting the essential role of these genes – and of ISGylation in general – in the IFNβ-mediated rescue of viability upon *Brca2* loss.

**Figure 3.**
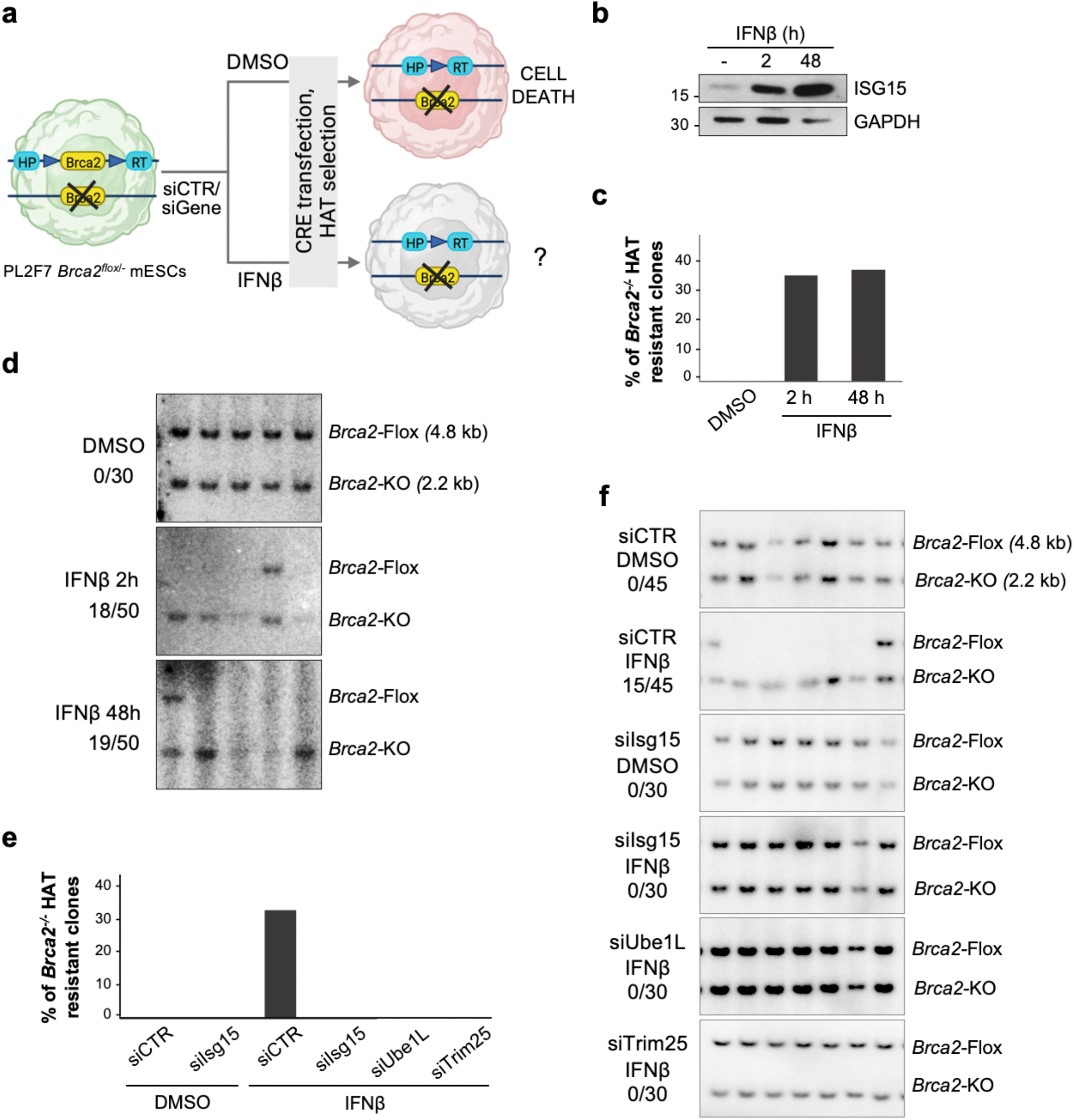
Upregulation of IFN/ISG15 restores viability in BRCA2-deficient mESCs. (**a**) Schematic for testing conditions for generation of *Brca2*^-/-^ in PL2F7. (**b**) ISG15 protein expression in PL2F7 cells after treatment with ± IFNβ (30 U/mL) either for 48 h in continuous or for 2 h pulse and 46 h chase. (**c,d**) Percentage of *Brca2*^-/-^ HAT resistant clones and representative Southern blot showing *Brca2*^*flox*/-^ or *Brca2*^-/-^ mESC upon ± IFNβ pre-treatment as in **b**. (**e,f**) Ratio of number of rescued clones and total numbers of HAT resistant clones analysed, and representative Southern blot showing *Brca2*^flox/-^ or *Brca2*^-/-^ mESC upon ± IFNβ (30 U/mL, 2 h) pre-treatment along with depletion of ISG15, UBE1L or TRIM25 by siRNA treatment.

### ISG15 is upregulated in BRCA1/2-deficient cells and is required for their fitness

Several lines of evidence indicate that conditions of replication fork instability and DNA damage lead to the accumulation of DNA fragments in the cytosol, which activate type I IFN via the cGAS/STING pathway (Lin and Pasero, 2021). As IFNβ strongly induces ISG15 expression, we reasoned that in genetic backgrounds characterized by genomic instability due to replication stress and DNA repair defects (such as conditions of BRCA1/2 deficiency), ISG15 expression might be elevated. Therefore, we compared ISG15 protein levels in isogenic pairs of BRCA1/2-deficient and -proficient cancer cells and found a consistent upregulation of ISG15 in cells carrying defects in either *BRCA1* or *BRCA2* (Figure 4a), suggesting that it may be required for the fitness of BRCA1/2-deficient cells. Indeed, ISG15 depletion markedly reduce the viability of MDA-MB-436 cells, which carry mutations in *BRCA1* gene, while MDA-MB-436 cells reconstituted with wild type *BRCA1* (Johnson et al., 2013) show only limited growth defects (Figure 4b and 4c).

**Figure 4.**
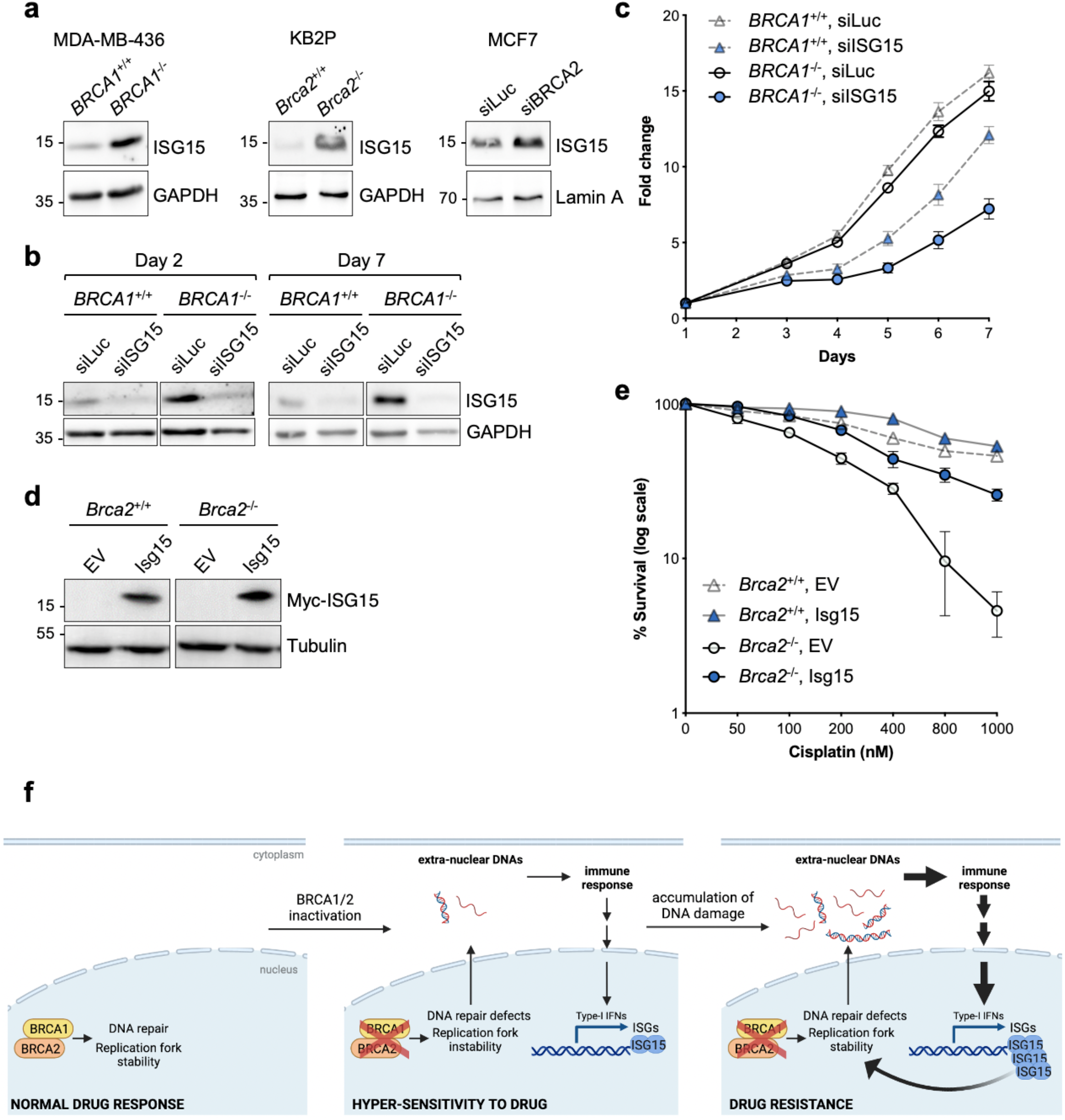
ISG15 is upregulated in BRCA1/2-deficient cells and is required for their fitness and reduced drug sensitivity. (**a**) Immunoblot showing ISG15 protein levels in MDA-MB-436, KB2P and MCF7 cells upon BRCA1 or BRCA2 depletion. (**b**) Immunoblot showing ISG15 protein levels in BRCA1-proficient or -deficient MDA-MB-436 cells at day 2 and day 7 post siISG15 treatment. GAPDH, loading control. (**c**) Cell proliferation graph showing fold change of viable cells (mean with SD) at indicated days normalized to cells at day 1 after seeding (n=3). (**d**) Immunoblot showing ISG15 protein levels in KB2P cells corresponding to Figure 4E. Tubulin, loading control. (**e**) Graph showing the percentage of surviving clones upon treatment with cisplatin with indicated doses in BRCA2-proficient (*Brca2*^+/+^ *p53*^-/-^) and -deficient (*Brca2*^-/-^ *p53*^-/-^) KB2P mouse cells upon ISG15 overexpression (n=3; error bars represent mean ± SEM). (**f**) Model depicting the contribution of IFN/ISG15 to drug response. Normal cells, characterized by efficient DNA repair and replication fork stability, show no sensitivity to chemotherapeutic drugs. Following the inactivation of DNA repair/replication factors, such as BRCA1/2, DNA repair and replication processes are impaired, resulting in genomic instability and the release of nucleic acids species into the cytosol. Over time, the accumulation of extra-nuclear DNAs leads to massive activation of the intracellular immune response, ultimately fostering a strong induction of the IFN-stimulated genes (ISGs), including ISG15 and its conjugation system (not shown), which restores stability of replication fork and favors the acquisition of drug resistance.

### Upregulation of ISG15 confers chemo-resistance to BRCA2-deficient cancer cells

Alterations in *BRCA1/2* genes are associated with marked genomic instability and cancer development. On the other hand, these conditions provide a window of opportunity for therapeutic intervention, since these tumor cells are exquisitely sensitive to chemotherapeutic drugs, such as cisplatin and PARP1 inhibitors (Bryant et al., 2005; Farmer et al., 2005; Fong et al., 2009). Unfortunately, BRCA1/2-deficient cancer cells very often acquire drug resistance, by means of different mechanisms, including reversion of *BRCA1/2* mutations, restoration of HR and of replication fork stability (Lord and Ashworth, 2013). As we observed that upregulation of ISG15 results in restoration of fork protection in BRCA1/2-deficient contexts, we asked whether it could also reduce sensitivity of these cells to cisplatin. To this purpose, we took advantage of the BRCA2-deficient KB2P cell line and its isogenic counterpart where the cell line has been reconstituted with human BRCA2. These cells were further engineered to obtain doxycycline-inducible expression of Myc-tagged mouse ISG15 (Figure 4d). In line with our previous results, IFNβ-mediated ISG15 induction restores protection of stalled forks in KB2P cells, although only partially (Figure S4a). Remarkably, cells defective in BRCA2 (*Brca2*^-/-^) show high sensitivity to cisplatin as compared to BRCA2-proficient cells, but upregulation of ISG15 dramatically reduces cisplatin sensitivity (Figure 4e and S4b).

## Discussion

It is well established that replication fork instability leads to accumulation of cytosolic nucleic acids, mimicking pathogen infection and therefore triggering the IFN-mediated immune response and inflammation. What is still unclear is how this inflammatory response in turn affects different aspects of DNA metabolism. Our work aims to close the circle and investigate how this IFN-mediated inflammatory response regulates DNA replication and repair processes, possibly impacting on cancer cell fitness and therapy response.

Tumors deficient in BRCA1/2 are associated with marked genomic instability, due to their pivotal functions in DNA repair via HR and DNA replication fork stability, which make them extremely sensitive to chemotherapeutic agents. Unfortunately, most patients develop drug resistance through different mechanisms, including restoration of HR and replication fork protection (Jaspers et al., 2013; Ray Chaudhuri et al., 2016). Here we show that activation of the IFN pathway restores the stability of nascent DNA in BRCA1/2-deficient contexts. Despite pleiotropic effects and numerous targets of IFN signaling, this effect is completely dependent on the ubiquitin-like protein ISG15 and on its conjugation system, indicating a key role for protein ISGylation in replication fork dynamics. ISG15-mediated fork protection is beneficial to cancer cells; indeed, the sole upregulation of ISG15 increases resistance of BRCA2-deficient mouse breast cancer cells to platinum-derived chemotherapeutic agents. These findings offer an explanation for the controversial data on the effect of type I IFN in cancer therapy. It is well established that the efficacy of several therapeutic strategies against cancer, including cytotoxic drugs, radiotherapy and targeted immunotherapies, depends on type I IFN signaling (Zitvogel et al., 2015). On the other hand, several studies indicate that IFN signaling may induce resistance to DNA damage and radiotherapy (Budhwani et al., 2018) and identified an IFN-related DNA damage resistance signature (IRDS) – including IFN response genes and ISG15 itself – correlating with acquired chemo- and radio-resistance (Khodarev et al., 2004; Weichselbaum et al., 2008). Our data show that IFN-induced ISG15 can promote drug resistance in specific genetic contexts, i.e., BRCAness, through the stabilization of nascent DNA strands. Intriguingly, while ISG15-mediated restoration of fork protection has been consistently observed in all cellular systems we have tested, the acquisition of drug resistance appears to be rather tumor and cell type-specific, probably depending on the complexity of the genetic background. This variability is likely to reflect differential expression of key factors, which are positively or negatively regulated by ISG15 and ISGylating enzymes. Future investigations are needed to identify those factors that, in combination with the upregulation of ISG15, are required to alter drug sensitivity of BRCA1/2-mutated cancer cells.

Another unexpected observation derives from the experiments in mESC showing that the IFN/ISG15 system restores viability of BRCA*2*-deficient cells. Importantly, also in this system we observed that IFNβ treatment protects stalled forks from degradation but does not rescue RAD51 *foci* formation. This result suggests that, in the absence of HR, the restoration of replication fork protection supports the viability of BRCA*2*-defective mESCs. Similarly, previous studies showed that suppression of fork degradation and reversed fork protection (as upon depletion of PTIP) promote ESC viability without impacting on RAD51 *foci* (Ding et al., 2016; Mijic et al., 2017; Ray Chaudhuri et al., 2016). In line with those observations, our data support the concept that defects in fork metabolism contribute to ESC lethality upon BRCA2 loss.

Notably, we observed that deficiency in *BRCA* genes increases *ISG15* expression, as compared to their syngeneic BRCA1/2-proficient counterparts, likely because of the accumulation of cytosolic DNA fragments and activation of the immune response. ISG15 upregulation is thus consistent and beneficial for these cells, as its depletion dramatically reduces cell viability. Over time, sustained ISG15 expression – fostered by BRCA deficiency – may *per se* lead to restoration of fork protection and promote drug resistance, suggesting the intriguing hypothesis that BRCAness conditions, and fork instability in general, are intrinsically prone to develop chemoresistance (Figure 4f). It will be important to extend these intriguing observations to other HR deficiencies.

Taken together, our results identify the IFN/ISG15 pathway as a key modulator of DNA replication fork protection and help explain the complex role of type I IFN – and ISG15 itself – in cancer development and drug response. Moreover, our data implicate that the IFN/ISG15 system should be carefully considered as a target to potentiate classical chemotherapy or as a critical modulator for immunotherapeutic options, especially when combined with DNA replication interference.

## Materials and methods

### Cell lines and cell culture

MEFs, U2OS and MCF7 cells were cultured in DMEM supplemented with 10% FBS and 0.05 U penicillin/streptomycin. U2OS Flp-In TREx cells were cultured in DMEM supplemented with 10% FBS, 0.05 U penicillin/streptomycin, 10 μg/mL blasticidin and 100 μg/mL hygromycin B. SUM149PT cells were cultured in adDMEM/F12 with 5% FBS, 0.05 U penicillin/streptomycin, 5 μg/mL of insulin, 10 mM HEPES, 2 mM L-Glutamine, 1 μg/mL of hydrocortisone. MDA-MB 436 *BRCA1*^-/-^ and *BRCA1*^+/+^ (reconstituted) cells were kindly gifted by Neil Johnson. Cells were maintained in RPMI media supplemented with 10% FBS, 0.05 U penicillin/streptomycin. CAPAN-1 cells were cultured in DMEM supplemented with 20% FBS, 0.05 U penicillin/streptomycin.

*Brca2*^+/+^ and *Brca2*^-/-^ mouse mammary tumor cells (KB2P 1.21) have been previously described (Jonkers et al., 2001). Both *Brca2*^+/+^ and *Brca2*^-/-^ cell lines were cultured under low oxygen conditions (3% O_2_, 5% CO_2_, 37°C) using DMEM (GIBCO; 41966-029) supplemented with 10% FBS, 50 U/mL penicillin, 50 μg/mL streptomycin, 5 μg/mL insulin (Sigma; I0516), 5 ng/mL murine epidermal growth factor (Sigma; E4127), and 5 ng/mL cholera toxin (List biological laboratories; 9100B).

All mouse embryonic stem cells (mESCs) were cultured on top of mitotically inactive STO-Neomycin-LIF-Puromycin (SNLP) feeder cells in M15 media [knockout DMEM media (Life Technologies) supplemented with 15% FBS (GE Life Sciences-Hyclone), 0.00072% β-mercaptoethanol, penicillin (100 U/mL), streptomycin (100 μg/mL), and L-glutamine (0.292 mg/ml)] at 37°C and 5% CO_2_. PL2F7 cells expressing BRCA2 R2336H variants were generated by electroporating respective bacterial artificial chromosomes in PL2F7 cells (Biswas et al., 2011; Kuznetsov et al., 2008). PL2F7-*Brca2*^-/-^;*BRCA2*(*R2336H*) cells only express the hypomorphic allele.

*Trim25*^+/+^ and *Trim25*^-/-^ MEFs were kindly gifted by Satoshi Inoue (Tokyo Metropolitan Institute of Gerontology, University of Tokyo, Japan). *Ube1L*^+/+^ and *Ube1L*^-/-^ MEFs were kindly gifted by Dong-Er Zhang (Moores Cancer Center, University of California, San Diego, USA). *Isg15*^+/+^ and *Isg15*^-/-^ MEFs were kindly gifted by Klaus-Peter Knobeloch (Institute of Neuropathology, University Clinic Freiburg, Germany).

### Lentiviral transduction

For lentivirus production, HEK293T cells were transfected with the packaging plasmids pVSV, pMDL, pREV and the expression plasmid pCW57.1-TRE-Myc-ISG15-IRES-EGFP or pCW57.1-TRE-IRES-EGFP (derived from pCW57.1-TRE kindly provided by Arnab Ray Chaudhuri) using jetPRIME^®^ (polyplus transfection) according to manufacturer’s instructions. The following day, medium was replaced with fresh DMEM medium, lentivirus was collected 72 h after transfection and stored at −80°C. Lentiviral titer was determined using FACS analysis (BD LSR II Fortessa) for GFP positive cells 72 h after transduction and doxycyclin treatment (1 μg/mL). For stable cell line generation, SUM149PT cells grown on 6-well plates were transduced overnight with lentiviral particles (MOI=1) in adDMEM/F12 supplemented with 10 μg/mL polybrene. Cells were washed 18 h after transduction and selected using Blasticidin antibiotics.

### DNA fiber assay

Following the depletion of proteins of interest or IFNβ treatment, cells were sequentially pulse-labeled with 33 μM CldU (Sigma-Aldrich; C6891) and 339 μM IdU (Sigma-Aldrich; I7125) for 20 min, or in case of U2OS 30 min. Following, the cells were treated with 4 mM HU (Sigma-Aldrich; H8627) for 4 h at 37°C. Cells were collected and resuspended in cold PBS to a concentration of 2.5×10^5^ labeled and 3.5×10^5^ unlabeled cells per mL. Labeled cells were mixed 1:2 (v/v) with unlabeled cells and 4 μL cells were lysed for 9 min with 7.5 μL lysis buffer (200 nM T ris-HCl pH 7.4, 50 mM EDTA, 0.5% (w/v) SDS) directly on a glass slide. Slides were tilted at 30-45° to stretch the DNA fibers, air-dried, and fixed in 3:1 methanol/acetic acid overnight at 4°C. The fibers were denatured with 2.5 M HCl for 1.5 h, washed with PBS and blocked for 40 min with 2% BSA/PBS-Tween. The CldU and IdU tracks were stained for 2.5 h with anti-BrdU primary antibodies recognizing CldU (1:500; Abcam; ab6326) and IdU (1:100; BD Biosciences; 347580), and for 1 h with secondary antibodies anti-mouse Alexa Fluor 488 (Invitrogen; A11001) and anti-rat Cy3 (Jackson Immuno Research; JAC712-166-153) in the dark. Coverslips were mounted using ProLong Gold Antifade Mountant. Images were acquired on a Leica DM6 B microscope at a lens-magnification of 63x and analyzed using ImageJ software (NIH). The pixel values were converted to μm (1 pixel corresponds to 0.146 μm) and IdU/CldU ratios were calculated as a measure of stalled replication fork degradation. Per experiment, at least 100 individual fibers were scored. Data were pooled from independent experiments. The number of biological replicates is indicated in the figure legend. Statistical differences in IdU/CldU ratios were determined by Kruskal-Wallis or Mann-Whitney test (GraphPad Prism 9).

### siRNA transfection

Cells were plated and transfected the following day for 48 or 72 h (as indicated below) using Oligofectamine transfection reagent (Invitrogen) according to manufacturer’s protocol Human cell lines were transfected with the following siRNAs: siLuc (5’-CGUACGCGGAAUACUUCGAUUdTdT-3’); siISG15 (50 nM, 72h; 5’-GCAACGAAUUCCAGGUGUCdTdT-3’); siUBE1L (50 nM, 72h; 5’-UAGUGCUGGCGUCUCAGCUUCUCCUdTdT-3’); siTRIM25 (50 nM, 72h; 5’-GGCUCAGAACACUUGAUAUTT-3’); siBRCA1 (40 nM, 48h; 5’-GGAACCUGUCUCCACAAAGTT-3’); siBRCA2 (40 nM, 48h; 5’-UUGACUGAGGCUUGCUCAGUUdTdT-3’); MEFs were transfected with a mix of the following siRNAs: siBrca2#1 (60 nM, 48h; 5’-UGUUAGGAGAUUCAUCUGGdTdT-3’); siBrca2#2 (60 nM, 48h; 5’-GGCCUAGUCUCAAGAACUCdTdT-3’); siBrca2#3 (60 nM, 48h; 5’-GGAAUUGUAAGGUAGGCUCdTdT-3’) mESC were transfected with the following siRNAs from Horizon Discovery: siIsg15 (25 nM, Cat. L-167630-00-0005) siUbe1L (25 nM, Cat. L-040733-01-0005) siTrim25 (25 nM, Cat. L-065539-01-0005)

### RNA extraction and cDNA preparation

Total RNA was extracted from cell pellets using TRIzol™ Reagent (Thermo Fischer; 15596026) according to the manufacturer’s protocol. RNA concentration was determined using a Thermo Scientific NanoDrop One. The total RNA was next reverse-transcribed to cDNA by M-MLV reverse transcriptase (Promega, M1701) 0.5 mg/mL Oligo(dT)_15_ primers were added to 200 ng of RNA and was incubated at 70°C for 5 mins. After 5 mins on ice, MMLV reverse-transcriptase master mix consisting of 5x Promega buffer, dNTP mix and M-MLV reverse transcriptase was added and incubated at 40°C for 30 min to synthesize the cDNA followed by 30 min incubation at 50°C to enhance synthesis from RNA with secondary structures. Finally, reverse transcriptase was inactivated by incubating at 70°C for 15 mins.

### qPCR

For qPCR the LightCycler^®^ SYBR Green system (Roche, 04707516001) was used. cDNA (11.42 ng) was mixed with 2x SYBR Green Master mix and 1 mM forward and reverse primers. The qPCR was run on a LightCycler^®^ 480 II (Roche).

TaqMan gene expression assays (Thermo Fisher Scientific) for *Trim25*^-/-^ and *Ube1L*^-/-^ MEFs were performed: *Brca2* (Mm00464783_m1), and *Hprt* (Mm03024075_m1). The samples were analyzed on a LightCycler^®^ 480 II instrument (Roche) and target mRNA abundance was subsequently calculated relative to *Hprt*.

### qPCR primers

Mouse *Isg15*

forward: 5’-GTGGTACAGAACTGCAGCGA-3’

reverse: 5’-TCAGCCAGAACTGGTCTTCG-3’

Mouse *Ube1L*

forward: 5’-GGAGTTAGGGCGAATGGAGG -3’

reverse: 5’ -GGAGTTAGGGCGAATGGAGG -3’

Mouse *Trim25*

forward: 5’-ATGGCTCAGGTAACAAGGGAG -3’

reverse: 5’ -GGGAGCAACAGGGGTTTTCTT -3’

Mouse *Brca2*

forward:5’-AGCCCAGCTTGAAGCAAGT -3’

reverse:5’ -GGATCATTCGGTAAACAGCG -3’

Mouse *Actin*

forward: 5’-CTGTCCCTGTATGCCTCTG -3’

reverse: 5’ -ATGTCACGCACGATTTCC -3’

Human *UBE1L*

forward: 5’-TGATGTTTGAGAAGGATGATG -3’

reverse: 5’ -CCGGTGGAATCCCGTAGTT -3’

Human *GAPDH*

forward: 5’-ACAACTTTGGTATCGTGGAAG -3’

reverse: 5’ -GCCATCACGCCACAGTTTC -3’

### Drugs and treatments

HU (Sigma-Aldrich; H8627) was dissolved in ddH2O at a 0.1 M concentration (7.6 mg/mL) and dissolved in growth medium to a final concentration of 4 mM.

Recombinant human IFNβ (PeproTech; 300-02BC) was diluted in growth medium to a concentration of 1000 U/mL and dissolved in growth medium to a final concentration of 30 U/mL.

Mouse IFNβ (Sigma-Aldrich; I9032-1VL) was diluted in growth medium to a concentration of 25’600 U/mL and was dissolved in growth medium to a final concentration of 30 U/mL.

Etoposide (Sigma-Aldrich; E1383) was dissolved in DMSO at a concentration of 10 mM and dissolved in growth medium to a final concentration of 5 μM.

Cisplatin (Sigma-Aldrich; 232120) was dissolved in PBS at a concentration of 10 mM and dissolved in growth medium to a final indicated concentration.

### Western blotting

Cells were lysed in RIPA buffer (50 mM Tris-HCl pH 7.4, 150 mM NaCl, 1% Triton X-100, 1% sodium deoxycholate, 0.1% SDS) or NP40 buffer (100 mM Tris-HCl pH 7.4, 300 mM NaCl, 2% NP40) supplemented with the following inhibitors: 50 mM sodium fluoride, 20 mM sodium pyrophosphate, 1 mM sodium orthovanadate, 1 mM phenylmethylsulfonyl fluoride (PMSF) and 1x Protease Inhibitor Cocktail (Sigma-Aldrich; P8340) for 10 min on ice. For total extracts, cells were treated with preheated (95°C) 1% SDS (10 mM Tris-HCl, pH 8.0; 1% SDS; 1 mM Na-Orthovanadate; 1x Protease Inhibitor Cocktail (Sigma-Aldrich; P8340)) and incubated for 10 min at 95°C. The lysates were sonicated by Bioruptor (Diagenode) at 4°C on the highest setting for 10 min (30s on and 30s off cycles) and centrifuged at 14’000 rpm for 10 min. Protein concentration was measured by Bradford protein assay (Bio-Rad; 5000006). Lysates were diluted with Laemmli Buffer (60 mM Tris-HCl pH 6.8, 2% SDS, 10% glycerol, 5% β-mercaptoethanol, 0.01% bromophenol blue), boiled at 95°C for 3 min or at 55°C for 10 min when BRCA2 was detected, equal amounts were loaded to polyacrylamide gels and ran at 160 V at room temperature. Proteins were blotted for 80 min (350 mA, room temperature) onto Amersham Protran 0.2 μm nitrocellulose membranes (GE Healthcare). Membranes were blocked in 5% milk in TBS-Tween for at least 2 h and incubated with primary antibodies overnight at 4°C. Following three washes in TBS-Tween, secondary antibodies were added for 1 h at room temperature. Membranes were washed three times in TBS-Tween and detected with WesternBright ECL HRP substrate (Advansta; K-12045-D50).

### List of antibodies used

1 List of antibodies used
| Antibody | Source | Cat | RRID | Dilution |
| --- | --- | --- | --- | --- |
| BRCA1 | Santa Cruz Biotechnology | sc-6954 | AB_626761 | 1:50 |
| BRCA2 | Millipore | OP95-100ug | AB_213443 | 1:500 |
| GAPDH | Millipore | MAB374 | AB_2107445 | 1:50'000 |
| ISG15 | K.P. Knobeloch Institute of Neuropathology, Freiburg, Germany | N/A | N/A | 1:1000 |
| ISG15 | Santa Cruz Biotechnology | sc-166755 | AB_2126308 | 1:1000 |
| Lamin A | Sigma-Aldrich | L1293 | AB_532254 | 1:1000 |
| Myc | Santa Cruz Biotechnology | sc-40 | AB_627268 | 1:1000 |
| PCNA | Santa Cruz Biotechnology | sc-56 | AB_628110 | 1:1000 |
| TRIM25 | Abcam | ab167154 | AB_2721902 | 1:2000 |
| Vinculin | Thermo Fisher Scientific | 700062 | AB_2532280 | 1:1000 |

### Immunofluorescence

Following siRNA transfection and IFNβ treatment, cells were grown on sterile 13-mm diameter glass coverslips. 48 h after siRNA transfection, cells were treated with 5 μM etoposide for 1 h or irradiated with 4 Gray on a Faxitron as indicated. Cells were washed with PBS, fixed with 4% paraformaldehyde for 10 min and permeabilized with 0.3% Triton X-100 in PBS for 5 min. Cells were blocked in 5% BSA in PBS for 1 h followed by incubation with primary antibodies for 1 h at room temperature. The following primary antibodies and dilutions were used: γH2AX (Millipore 05-635; 1:800), RAD51 (Bioacademia 70-001; 1:2000). Cells were washed with PBS and incubated with secondary antibodies for 30 min at room temperature. Total DNA was stained with 4’,6-diamidino-2-phenylidole (DAPI, 1 μM) for 5 min at room temperature. Images were acquired on a Leica DM6 microscope.

### Quantitative image-based cytometry (QIBC)

Images were acquired using Leica DM6 microscope under non-saturating conditions and identical settings were applied to all samples within one experiment. Images were then converted to file system suitable for analysis with Olympus ScanR Image analysis software (version 3.0.1). Nuclei segmentation was performed using DAPI signal, and further RAD51 foci detection performed using integrated spot detection module. The quantification of RAD51 foci were exported and analyzed using Spotfire data visualization software (TIBCO, version 7.0.1). Quantifications of three independent experiments are shown.

### Cell proliferation assay

300’000 cells were plated in 6 cm plates 24 h prior to IFNβ stimulation. 30 U/mL of IFNβ was added and incubated for 2 h and then washed and incubated with culture medium. 24 h after IFNβ induction cells were trypsinized and 10’000 cells were plated in each well of a 12 wells plate. Cell Titer Blue assay (Promega, G8080) was performed to access cell viability according to manufacturer’s protocol. Relative proliferation rate was quantified and normalized to day 1 (48 h post IFNβ induction).

### FACS

BRCA2-proficient (*Brca2*^+/+^ *p53*^-/-^) cell lines and, BRCA2-deficient (*Brca2*^-/-^ *p53*^-/-^) KB2P cell lines derived from mouse mammary tumor were transduced by lentiviral transduction. Post transduction, cells were selected with Blasticidin (10 μg/mL) for ten days and then EV and ISG15 expression were induced by 48 h of doxycycline (2 μg/mL). Transduced cells carrying the GFP construct were sorted by FACS (BD FACSAria™ III Cell Sorter) with a 488 nm argon ion laser based on their GFP fluorescence (using BD FACSDiva 9.0.1 software). Sorted GFP positive cells were kept in culture for 1 week and then used for clonogenic assay.

### Clonogenic assay

Clonogenic assay was performed according to the standard protocol as previously mentioned (Mukherjee et al., 2019). EV and ISG15 were induced by 48 h of doxycycline (2 μg/mL) in pCW57.1 GFP and pCW57.1 Myc-ISG15 WT-GFP in the *Brca2*^+/+^ and *Brca2*^-/-^ mouse KB2P transduced cell lines. GFP-positive cells were sorted by FACS and used for colony survival assay.

Cells were seeded in 6 cm cell culture dishes at low density with and without doxycycline (2 μg/mL). Twenty-four hours after seeding, cells were treated with Cisplatin at different concentrations for 6 h. Post-treatment, drug treated medium was washed out and cells were allowed to grow in a complete growth medium for 7 days in the presence and absence of doxycycline. The colonies detected were fixed, stained with Brilliant Blue R (Sigma-Aldrich; B0149) and subsequently analyzed with the Gel-counter by Oxford Optronix and appertaining Software (version 1.1.2.0). The survival was plotted after combining three independent experiments as the mean surviving percentage of colonies after drug treatment compared to the mean surviving colonies from the non-treated samples.

### Embryonic stem cell viability assay

PL2F7 cells were used for cell viability rescue experiments in *Brca2* conditional knockout ESCs. Viability assay was performed as described in (Kuznetsov et al., 2008).

### Southern blot

EcoRV-digested DNA was electrophoresed on 1% agarose gel in 1X TBE (0.1 M Tris, 0.1 M boric acid and 2 mM EDTA, pH 8.0) and transferred to nylon membrane. DNA probe for distinguishing conditional *Brca2* allele (*Brca2*-Flox, 4.8 kb) and *Brca2* knockout allele (*Brca2*-KO, 2.2 kb) 20 was labelled by [a-32P]-dCTP by Prime-It II Random Primer Labeling Kit (Agilent Technologies) and hybridized with Hybond-N. nylon membrane (GE Healthcare) at 65°C overnight. Membrane was washed twice with saline sodium citrate phosphate (SSCP) buffer containing 0.1% SDS in and exposed in phosphor image screen overnight and subsequently developed in Typhoon image scanner.

### Statistics

Number of biological replicates is defined in the legends of the figures. Results were analyzed using GraphPad, using Kruskal-Wallis or Mann-Whitney test, (two tailed P value. P value > 0.05 was considered not significant–ns-). The RAD51 *foci* counts were extracted from the raw data and subjected to statistical analysis using GraphPad Prism 9 (two tailed P value). The results were analyzed using Spotfire and GraphPad Prism9 using Kruskal-Wallis test. In cell proliferation assay, absorbance of each sample (technical triplicate) was normalized on untreated samples.

## Availability of data, materials and methods

Further information and requests for reagents and resources should be directed to and will be fulfilled by the Lead Contact, Lorenza Penengo.

All unique/stable reagents generated in this study will be made available upon request to the Lead Contact.

## Acknowledgements

We thank Sven Rottenberg, Dong-Er Zhang, Klaus-Peter Knobeloch, Satoshi Inoue and Neil Johnson for sharing reagents. This work was supported by the Intramural Research Program, Center for Cancer Research, National Cancer Institute, US National Institutes of Health to S.K.S., by NWO VIDI grant (VI.Vidi.193.131) and Dutch Cancer Society (KWF-11008) to A.R.C., by the Swiss Cancer Research foundation (KFS-4577-08-2018), the Worldwide Cancer Research (22-0181) and the Swiss National Science Foundation (SNSF; grants 310030_184966) to L.P.

## Author contributions

U.B. and R.N.M. performed the DNA fiber experiments, the immunofluorescence and proliferation studies; S.S.K. and S.K.S. designed and performed the viability studies in mESCs; P.D. and A.R.C. performed the drug sensitivity studies in mouse KB2P cells. L.P. conceived the project, designed the experiments, and wrote the manuscript, supported by U.B. and R.N.M.

## Supplementary figure legends

**Supplementary Figure 1.**
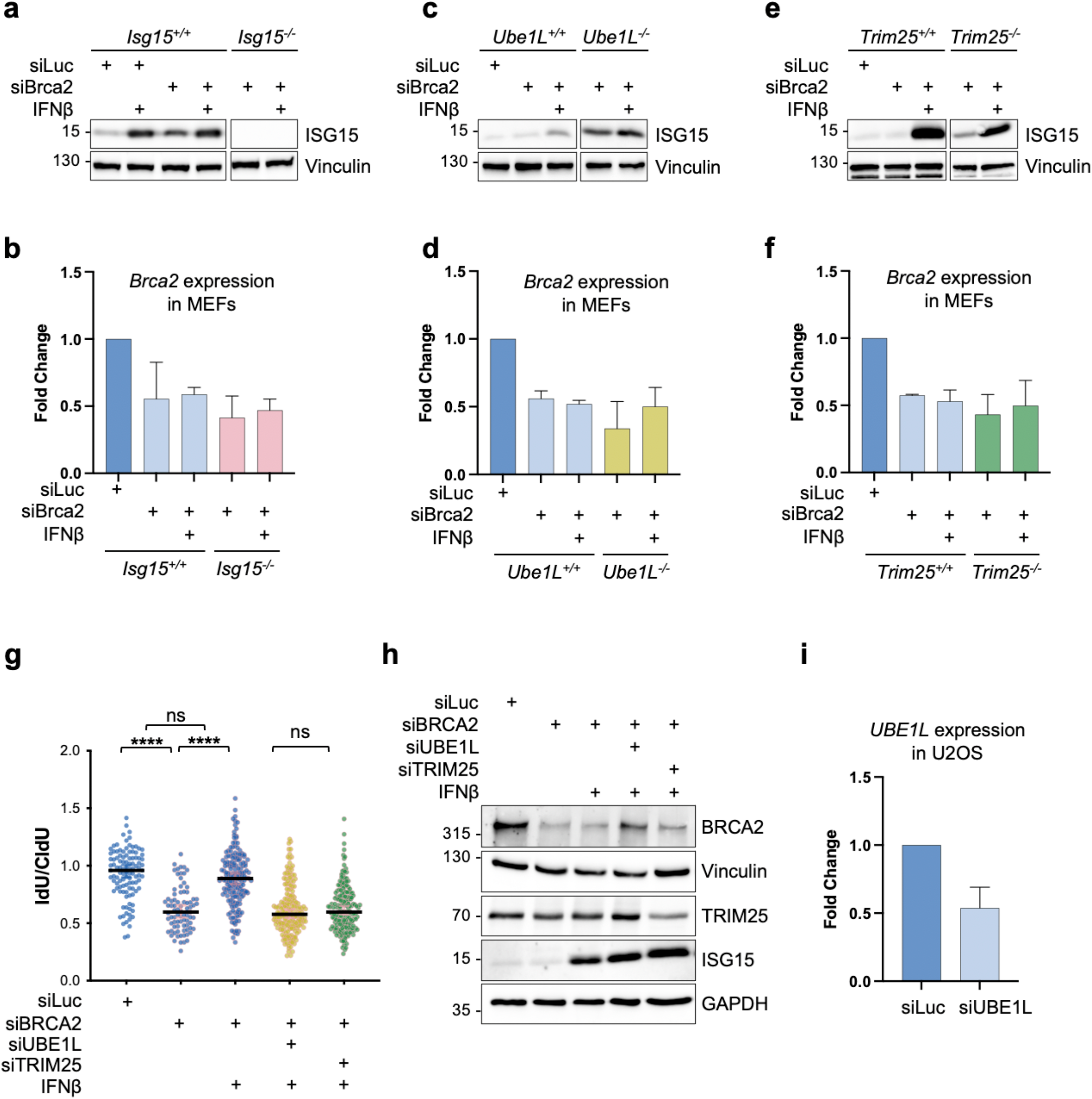
IFN/ISG15 restores fork protection in BRCA-deficient cells via ISG15 conjugation. (**a**) Immunoblot showing levels of ISG15 expression in parental and *Isg15*^-/-^ MEF cells with IFNβ treatment corresponding to Figure **2c**. Vinculin, loading control. (**b**) *Brca2* expression in parental and *Isg15*^-/-^ MEF cells measured by qPCR corresponding to Figure **2c**. (**c**) Immunoblot showing levels of ISG15 expression in parental and *Ube1L^-/-^* MEF cells with IFNβ treatment corresponding to Figure **2I**. Vinculin, loading control. (**d**) *Brca2* expression in parental and *Ube1L*^-/-^ MEF cells measured by qPCR corresponding to Figure **2i**. (**e**) Immunoblot showing levels of ISG15 expression in parental and *Trim25*^-/-^ MEF cells with IFNβ treatment corresponding to Figure **2j**. Vinculin, loading control. (**f**) *Brca2* expression in parental and *Trim25*^-/-^ MEF cells measured by qPCR corresponding to Figure **2j**. (**g**) IdU/CldU ratio analysis in siLuc, siBRCA2 or double depletion with siBRCA2 along with siUBE1L or siTRIM25 in U2OS cells treated with ± IFNβ (30 U/mL, 2 h) and chased for 46 h (n=2). Horizontal lines represent the median value. Statistical analysis according to Kruskal-Wallis test was performed; ****, P<0.0001. (**h**) Immunoblot showing levels of BRCA2, TRIM25 and ISG15 in U2OS cells with indicted siRNA treatment along with ± IFNβ (30 U/mL, 2 h) and chased for 46 h corresponding to Figure **S1g**. Vinculin and GAPDH, loading controls. (**i**) *UBE1L* expression in U2OS cells upon indicated siRNA treatment measured by qPCR corresponding to Figure **S1g**.

**Supplementary Figure 2.**
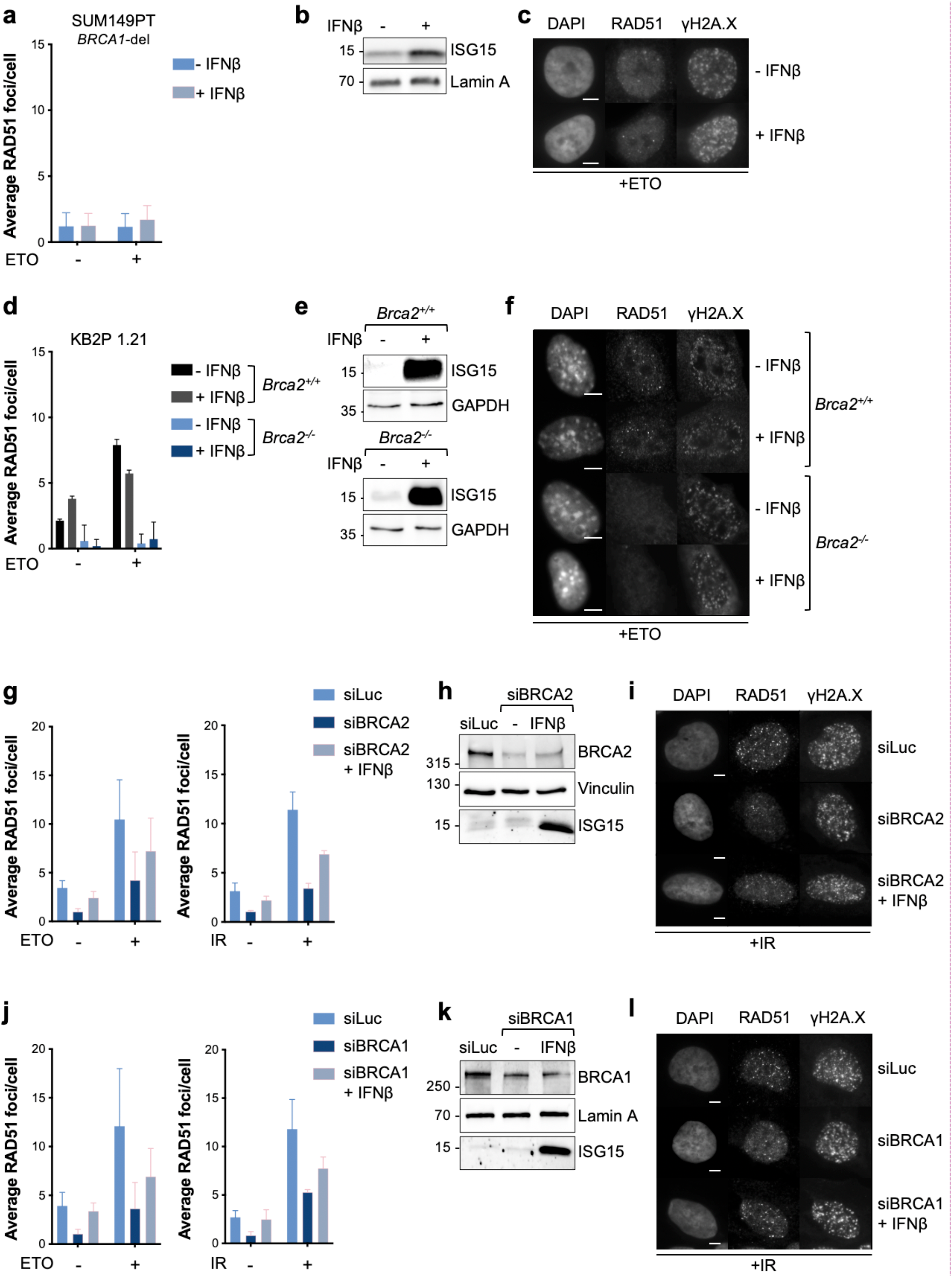
IFN/ISG15 does not restore homologous recombination in BRCA-deficient cells. (**a**) Average number of RAD51 *foci* per cell in SUM149PT cells upon ± IFNβ (30 U/mL, 2 h; 46 h chase). Cells were optionally treated with 5 μM etoposide (ETO) for 1 h and subsequently stained with antibodies against RAD51 and γH2AX. Data are represented as mean + SD (n=3). (**b**) ISG15 expression in untreated and IFNβ-treated cells as in **a**. Lamin A, loading control. (**c**) Representative images of RAD51 formation in SUM149PT cells treated with ETO upon ± IFNβ as in **a**. Scale bars, 5 μm. (**d**) Average number of RAD51 *foci* formation in KB2P cells (*Brca2*^+/+^ *p53*^-/-^ and *Brca2*^-/-^ *p53*^-/-^) treated with ± IFNβ (30 U/mL, 2h) and chased for 46h. Cells were treated with 5 μM of etoposide for 1 h and incubated with EdU for 30 min. Cells were fixed and subsequently stained with antibodies against RAD51 and γH2AX. Data are represented as mean + SD (n=3). (**e**) ISG15 expression in untreated and IFNβ-treated cells as in **d**. Gapdh, loading control. (**f**) Representative images of RAD51 *foci* formation in KB2P cells (*Brca2^+/+^* and *Brca2*^-/-^) treated with ETO upon ± IFNβ as in **d**. Scale bars, 5 μm. (**g**) Average number of RAD51 *foci* per cell in untreated, ETO-treated or irradiated (4 Gray) siBRCA2 or siLuc U2OS cells with ± IFNβ (30 U/mL, 2 h; 46 h chase). One hour after irradiation cells were fixed and subsequently stained with antibodies against RAD51 and γH2AX. Data are represented as mean + SD (n=3). (**h**) BRCA2 and ISG15 expression as in **g**. Vinculin, loading control. (**i**) Representative images of RAD51 *foci* formation in siBRCA2 or siLuc U2OS cells treated with ± IFNβ and IR as in **g**. Scale bars, 5 μm. (**j**) Average number of RAD51 *foci* per cell in untreated or irradiated (4 Gray) siBRCA1 or siLuc U2OS treated cells with ± IFNβ (30 U/mL, 2 h; 46 h chase). One hour after irradiation cells were fixed and subsequently stained with antibodies against RAD51 and γH2AX. Data are represented as mean + SD. (**k**) Representative images of RAD51 *foci* formation in siBRCA1 or siLuc U2OS cells treated with ± IFNβ as in **j**. Scale bars, 5 μm. (**l**) BRCA1 and ISG15 expression as in **j**. Lamin A, loading control.

**Supplementary Figure 3.**
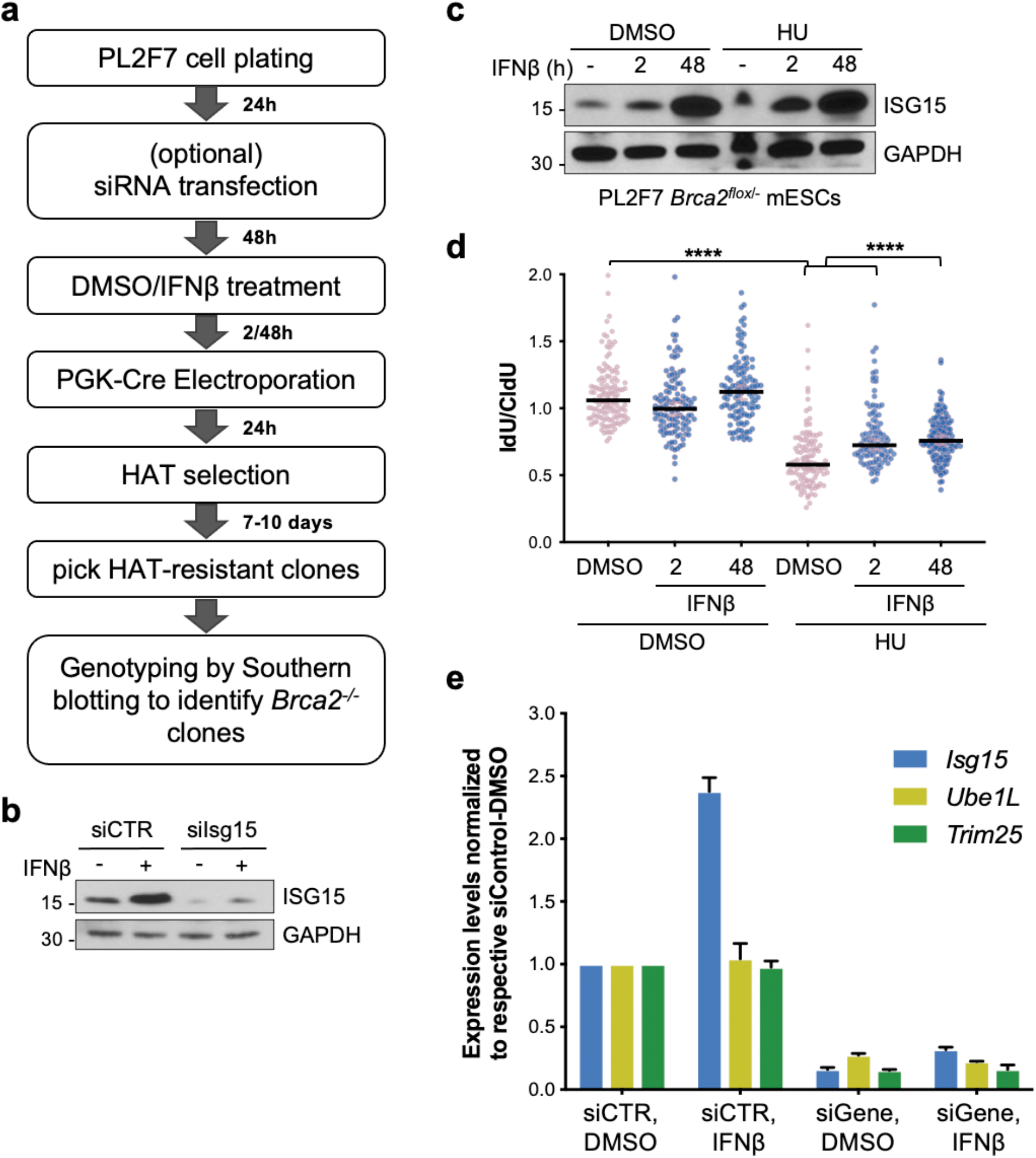
Upregulation of IFN/ISG15 restores viability in BRCA2-deficient mESCs. (**a**) Schematic representation of the experimental workflow to test the effect of the IFN system on the viability of *Brca2*^-/-^ mESCs. (**b**) Immunoblot showing the levels of ISG15 upon induction with ± IFNβ (30 U/mL, 2 h) pre-treatment and depletion by siRNA. GAPDH, loading control. (**c**) IdU/CldU ratio analysis in PL2F7-*Brca2*^-/-^;BRCA2 (R2336H) cells treated with ± IFNβ (30 U/mL, 2 h; 46 h chase). Statistical analysis was performed using Mann-Whitney test; ****, P<0.0001. (**d**) Immunoblot showing ISG15 protein levels upon IFNβ (30 U/mL, 2h) pre-treatment along with either DMSO or with hydroxyurea (HU, 4 mM) for 3 h corresponding to Figure **S3e**. (**e**) Expression levels of *Isg15*, *Ube1L* and *Trim25* in mESC with ± IFNβ (30 U/mL, 2 h) pre-treatment and depletion by indicated siRNA.

**Supplementary Figure 4.**
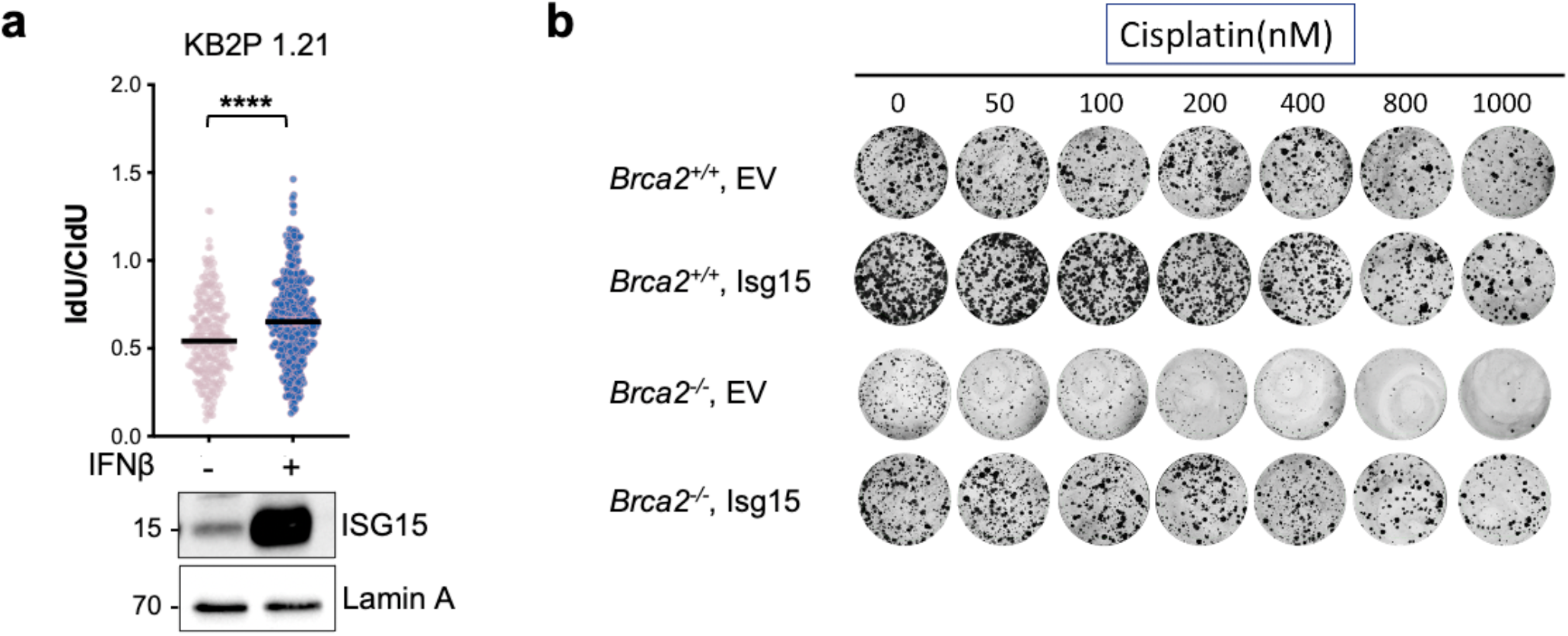
ISG15 up-regulation promotes fork protection, but not Rad51 *foci*, in KB2P cells and reduces drug sensitivity. (**a**) IdU/CldU ratio analysis in *Brca2*^-/-^ *p53*^-/-^ KB2P cells treated with ± IFNβ (30 U/mL, 2 h; 46 h chase) (n=3). Horizontal lines represent the median value. Statistical analysis according to Mann-Whitney test was performed; ns, non-significant; ****, P<0.0001. Lower panel showing ISG15 protein levels. (**b**) Representative colony images of the clonogenic assay upon increasing doses of Cisplatin.

## References

Bakhoum, S., Ngo, B., Bakhoum, A., Cavallo-Fleming, J.A., Murphy, C.W., Powell, S.N., and Cantley, L. (2018). Chromosomal Instability Drives Metastasis Through a Cytosolic DNA Response. Int. J. Radiat. Oncol. 102.

Biswas, K., Das, R., Alter, B.P., Kuznetsov, S.G., Stauffer, S., North, S.L., Burkett, S., Brody, L. C., Meyer, S., Byrd, R.A., et al. (2011). Acomprehensive functional characterization of BRCA2 variants associated with Fanconi anemia using mouse ES cell-based assay. Blood 118.

Bryant, H.E., Schultz, N., Thomas, H.D., Parker, K.M., Flower, D., Lopez, E., Kyle, S., Meuth, M., Curtin, N.J., and Helleday, T. (2005). Specific killing of BRCA2-deficient tumours with inhibitors of poly(ADP-ribose) polymerase. Nature 434, 913–917.

Budhwani, M., Mazzieri, R., and Dolcetti, R. (2018). Plasticity of Type I Interferon-Mediated Responses in Cancer Therapy: From Anti-tumor Immunity to Resistance. Front Oncol 8, 322.

Cavanagh, H., and Rogers, K.M. (2015). The role of BRCA1 and BRCA2 mutations in prostate, pancreatic and stomach cancers. Hered Cancer Clin Pr. 13, 16.

Ding, X., Chaudhuri, A.R., Callen, E., Pang, Y., Biswas, K., Klarmann, K.D., Martin, B.K., Burkett, S., Cleveland, L., Stauffer, S., et al. (2016). Synthetic viability by BRCA2 and PARP1/ARTD1 deficiencies. Nat. Commun. 7.

Erdal, E., Haider, S., Rehwinkel, J., Harris, A.L., and McHugh, P.J. (2017). A prosurvival DNA damage-induced cytoplasmic interferon response is mediated by end resection factors and is limited by Trex1. Genes Dev. 31.

Evers, B., Drost, R., Schut, E., de Bruin, M., van der Burg, E., Derksen, P.W., Holstege, H., Liu, X., van Drunen, E., Beverloo, H.B., et al. (2008). Selective inhibition of BRCA2-deficient mammary tumor cell growth by AZD2281 and cisplatin. Clin Cancer Res 14, 3916–3925.

Farmer, H., McCabe, N., Lord, C.J., Tutt, A.N., Johnson, D.A., Richardson, T.B., Santarosa, M., Dillon, K.J., Hickson, I., Knights, C., et al. (2005). Targeting the DNA repair defect in BRCA mutant cells as a therapeutic strategy. Nature 434, 917–921.

Fong, P.C., Boss, D.S., Yap, T.A., Tutt, A., Wu, P., Mergui-Roelvink, M., Mortimer, P., Swaisland, H., Lau, A., O’Connor, M.J., et al. (2009). Inhibition of poly(ADP-ribose) polymerase in tumors from BRCA mutation carriers. N Engl J Med 361, 123–134.

Gowen, L.C., Johnson, B.L., Latour, A.M., Sulik, K.K., and Koller, B.H. (1996). Brca1 deficiency results in early embryonic lethality characterized by neuroepithelial abnormalities. Nat. Genet. 12.

Hakem, R., De La Pompa, J.L., Sirard, C., Mo, R., Woo, M., Hakem, A., Wakeham, A., Potter, J., Reitmair, A., Billia, F., et al. (1996). The tumor suppressor gene Brca1 is required for embryonic cellular proliferation in the mouse. Cell 85.

Han, H.G., Moon, H.W., and Jeon, Y.J. (2018). ISG15 in cancer: Beyond ubiquitin-like protein. Cancer Lett 438, 52–62.

Harding, S.M., Benci, J.L., Irianto, J., Discher, D.E., Minn, A.J., and Greenberg, R.A. (2017). Mitotic progression following DNA damage enables pattern recognition within micronuclei. Nature 548.

Härtlova, A., Erttmann, S.F., Raffi, F.A.M., Schmalz, A.M., Resch, U., Anugula, S., Lienenklaus, S., Nilsson, L.M., Kröger, A., Nilsson, J.A., et al. (2015). DNA Damage Primes the Type I Interferon System via the Cytosolic DNA Sensor STING to Promote Anti-Microbial Innate Immunity. Immunity 42.

Jaspers, J.E., Kersbergen, A., Boon, U., Sol, W., van Deemter, L., Zander, S.A., Drost, R., Wientjens, E., Ji, J., Aly, A., et al. (2013). Loss of 53BP1 causes PARP inhibitor resistance in Brca1-mutated mouse mammary tumors. Cancer Discov 3, 68–81.

Johnson, N., Johnson, S.F., Yao, W., Li, Y.C., Choi, Y.E., Bernhardy, A.J., Wang, Y., Capelletti, M., Sarosiek, K.A., Moreau, L.A., et al. (2013). Stabilization of mutant BRCA1 protein confers PARP inhibitor and platinum resistance. Proc. Natl. Acad. Sci. U. S. A. 110.

Jonkers, J., Meuwissen, R., van der Gulden, H., Peterse, H., van der Valk, M., and Berns, A. (2001). Synergistic tumor suppressor activity of BRCA2 and p53 in a conditional mouse model for breast cancer. Nat Genet 29, 418–425.

Khodarev, N.N., Beckett, M., Labay, E., Darga, T., Roizman, B., and Weichselbaum, R.R. (2004). STAT1 is overexpressed in tumors selected for radioresistance and confers protection from radiation in transduced sensitive cells. Proc. Natl. Acad. Sci. U. S. A. 101.

Kuznetsov, S.G., Liu, P., and Sharan, S.K. (2008). Mouse embryonic stem cell-based functional assay to evaluate mutations in BRCA2. Nat. Med. 14.

Li, T., and Chen, Z.J. (2018). The cGAS-cGAMP-STI NG pathway connects DNA damage to inflammation, senescence, and cancer. J. Exp. Med. 215.

Lin, Y.L., and Pasero, P. (2021). Replication stress: from chromatin to immunity and beyond. Curr. Opin. Genet. Dev. 71.

Lord, C.J., and Ashworth, A. (2013). Mechanisms of resistance to therapies targeting BRCA-mutant cancers. Nat Med 19, 1381–1388.

MacKenzie, K.J., Carroll, P., Martin, C.A., Murina, O., Fluteau, A., Simpson, D.J., Olova, N., Sutcliffe, H., Rainger, J.K., Leitch, A., et al. (2017). CGAS surveillance of micronuclei links genome instability to innate immunity. Nature 548.

Mijic, S., Zellweger, R., Chappidi, N., Berti, M., Jacobs, K., Mutreja, K., Ursich, S., Ray Chaudhuri, A., Nussenzweig, A., Janscak, P., et al. (2017). Replication fork reversal triggers fork degradation in BRCA2-defective cells. Nat Commun 8, 859.

Mukherjee, C., Tripathi, V., Manolika, E.M., Heijink, A.M., Ricci, G., Merzouk, S., de Boer, H.R., Demmers, J., van Vugt, M.A.T.M., and Ray Chaudhuri, A. (2019). RIF1 promotes replication fork protection and efficient restart to maintain genome stability. Nat. Commun. 10.

Park, J.M., Yang, S.W., Yu, K.R., Ka, S.H., Lee, S.W., Seol, J.H., Jeon, Y.J., and Chung, C.H. (2014). Modification of PCNA by ISG15 plays a crucial role in termination of error-prone translesion DNA synthesis. Mol Cell 54, 626–638.

Parkes, E.E., Walker, S.M., Taggart, L.E., McCabe, N., Knight, L.A., Wilkinson, R., McCloskey, K.D., Buckley, N.E., Savage, K.I., Salto-Tellez, M., et al. (2017). Activation of STING-dependent innate immune signaling by s-phase-specific DNA damage in breast cancer. J. Natl. Cancer Inst. 109.

Perng, Y.C., and Lenschow, D.J. (2018). ISG15 in antiviral immunity and beyond. Nat. Rev. Microbiol. 16.

Raso, M.C., Djoric, N., Walser, F., Hess, S., Schmid, F.M., Burger, S., Knobeloch, K.P., and Penengo, L. (2020). Interferon-stimulated gene 15 accelerates replication fork progression inducing chromosomal breakage. J Cell Biol 219.

Ray Chaudhuri, A., Callen, E., Ding, X., Gogola, E., Duarte, A.A., Lee, J.E., Wong, N., Lafarga, V., Calvo, J.A., Panzarino, N.J., et al. (2016). Replication fork stability confers chemoresistance in BRCA-deficient cells. Nature 535, 382–387.

Rickman, K., and Smogorzewska, A. (2019). Advances in understanding DNA processing and protection at stalled replication forks. J. Cell Biol. 218.

Dos Santos, P.F., and Mansur, D.S. (2017). Beyond ISGlylation: Functions of Free Intracellular and Extracellular ISG15. J Interf. Cytokine Res 37, 246–253.

Schlacher, K., Christ, N., Siaud, N., Egashira, A., Wu, H., and Jasin, M. (2011). Double-strand break repair-independent role for BRCA2 in blocking stalled replication fork degradation by MRE11. Cell 145, 529–542.

Schlacher, K., Wu, H., and Jasin, M. (2012). A distinct replication fork protection pathway connects Fanconi anemia tumor suppressors to RAD51-BRCA1/2. Cancer Cell 22, 106–116.

Scully, R., Panday, A., Elango, R., and Willis, N.A. (2019). DNA double-strand break repair-pathway choice in somatic mammalian cells. Nat Rev Mol Cell Biol.

Sharan, S.K., Morimatsu, M., Albrecht, U., Lim, D.S., Regel, E., Dinh, C., Sands, A., Eichele, G., Hasty, P., and Bradley, A. (1997). Embryonic lethality and radiation hypersensitivity mediated by Rad51 in mice lacking Brca2. Nature 386.

Swaim, C.D., Scott, A.F., Canadeo, L.A., and Huibregtse, J.M. (2017). Extracellular ISG15 Signals Cytokine Secretion through the LFA-1 Integrin Receptor. Mol Cell 68, 581–590 e5.

Villarroya-Beltri, C., Guerra, S., and Sanchez-Madrid, F. (2017). ISGylation - a key to lock the cell gates for preventing the spread of threats. J Cell Sci 130, 2961–2969.

Weichselbaum, R.R., Ishwaran, H., Yoon, T., Nuyten, D.S., Baker, S.W., Khodarev, N., Su, A.W., Shaikh, A.Y., Roach, P., Kreike, B., et al. (2008). An interferon-related gene signature for DNA damage resistance is a predictive marker for chemotherapy and radiation for breast cancer. Proc Natl Acad Sci U S A 105, 18490–18495.

Wong, J.J., Pung, Y.F., Sze, N.S., and Chin, K.C. (2006). HERC5 is an IFN-induced HECT-type E3 protein ligase that mediates type I IFN-induced ISGylation of protein targets. Proc Natl Acad Sci U S A 103, 10735–10740.

Zitvogel, L., Galluzzi, L., Kepp, O., Smyth, M.J., and Kroemer, G. (2015). Type I interferons in anticancer immunity. Nat. Rev. Immunol. 15.

Zou, W., and Zhang, D.E. (2006). The interferon-inducible ubiquitin-protein isopeptide ligase (E3) EFP also functions as an ISG15 E3 ligase. J. Biol. Chem. 281.

